# Pathogen lifestyle determines host genetic signature of quantitative disease resistance loci in oilseed rape (*Brassica napus*)

**DOI:** 10.1101/2023.08.02.551671

**Authors:** Catherine N. Jacott, Henk-jan Schoonbeek, Gurpinder Singh Sidhu, Burkhard Steuernagel, Rachel Kirby, Xiaorong Zheng, Andreas von Tiedermann, Violetta K. Macioszek, Andrzej K. Kononowicz, Heather Fell, Bruce D.L. Fitt, Georgia K. Mitrousia, Henrik U. Stotz, Christopher J. Ridout, Rachel Wells

## Abstract

- Crops are affected by several pathogens, but these are rarely studied in parallel to identify common and unique genetic factors controlling diseases. Broad-spectrum quantitative disease resistance (QDR) is desirable for crop breeding as it confers resistance to several pathogen species.
- Here, we use associative transcriptomics (AT) to identify candidate gene loci associated with *Brassica napus* QDR to four contrasting fungal pathogens: *Alternaria brassicicola*, *Botrytis cinerea*, *Pyrenopeziza brassicae* and *Verticillium longisporum*.
- We did not identify any loci associated with broad-spectrum QDR to fungal pathogens with contrasting lifestyles. Instead, we observed QDR dependent on the lifestyle of the pathogen—hemibiotrophic and necrotrophic pathogens had distinct QDR responses and associated loci, including some loci associated with early immunity. Furthermore, we identify a genomic deletion associated with resistance to *V. longisporum* and potentially broad-spectrum QDR.
- This is the first time AT has been used for several pathosystems simultaneously to identify host genetic loci involved in broad-spectrum QDR.
- We highlight candidate loci for broad-spectrum QDR with no antagonistic effects on susceptibility to the other pathogens studies as candidates for crop breeding.

## Introduction

Crop-infecting fungi are responsible for approximately 20% of global crop losses annually (Fisher *et al*., 2018). Developing broad-spectrum resistance is vital for effective crop disease management, as it protects against several pathogen species or most races or isolates of the same pathogen (Kou and Wang, 2010). Despite this, plant interactions with different pathogens are rarely studied in collaborative programs to identify common and unique genetic factors controlling the diseases.

Most research and breeding focus on *‘R’* (resistance) gene-mediated qualitative resistance, often providing complete resistance to certain races of a pathogen (McDowell and Woffenden, 2003). *R* gene-mediated resistance can be effective at controlling disease but often breaks down when pathogen effector genes mutate. In contrast, quantitative disease resistance (QDR)—also known as partial resistance—presents a continuous distribution of phenotypes from susceptible to resistant.

QDR has great potential for crop improvement due to its broad-spectrum and durable nature; however, it is not but is not frequently targeted in breeding efforts due to a lack of insights into its underlying molecular determinants (Roux *et al*., 2014; Nelson *et al*., 2018). QDR is generally conferred by multiple genes, limiting the ability of pathogens to evolve resistance. QDR is likely broad-spectrum as it is often associated with constitutive features that affect the growth of diverse pathogens such as host plant morphology, basal defense, and signal transduction (Raman *et al*., 2020, Amas *et al*., 2021, Pliet-Nayel *et al*., 2017). Constitutive QDR may also function in combination with pathogen-specific resistance to prolong the effectiveness of *R*-genes (Pilet-Nayel *et al*., 2017, Brun *et al*., 2010). Although few QDR genes have been cloned and functionally validated, reported mechanisms include enhanced cell wall synthesis (Wisser *et al*., 2005; Corwin *et al*., 2016), enhanced secondary metabolism (Benson *et al*., 2015) modified transport processes (Moore *et al*., 2015; Deppe *et al*., 2018) hormone synthesis and transcriptional regulation (Qasim *et al*., 2020).

Identifying loci involved in broad-spectrum QDR to multiple pathogens is challenging. Different pathogens employ distinct infection and feeding strategies, including feeding on living host cells (biotrophy); killing plant cells to feed on their contents (necrotrophy); and keeping cells alive while establishing infection before switching to a necrotrophic mode (hemibiotrophy) (Kemen and Jones, 2012). These diverse strategies necessitate differing plant defense mechanisms for effective resistance. For example, reactive oxygen species (ROS) production and cell death can provide resistance to biotrophic and hemibiotrophic pathogens (Jia *et al*., 2000; Vleeshouwers *et al*., 2000, Vetter *et al*., 2012). However, ROS accumulation and cell death can promote pathogenicity for some necrotrophic pathogens that use necrosis to facilitate their colonization (Tiedemann, 1997; Govrin and Levine, 2000; Lorang *et al*., 2007; Williams et al., 2011, Faris and Friesen, 2020). Additionally, certain plant hormones salicylic acid (SA) and jasmonic acid (JA) exhibit opposing effects on resistance dependent on pathogen lifestyle (Navarro, *et al*., 2008): the activation of the JA-signaling provides resistance against necrotrophic pathogens; but the activation of SA-signaling protects against biotrophic and hemibiotrophic pathogens (Glazebrook, 2005).

Certain broad-spectrum QDR mechanisms could relate to innate immunity based on the detection of shared pathogenic features irrespective of their lifestyle. For example, some quantitative trait loci (QTLs) implicated in QDR resemble cell surface pattern recognition receptors (PRRs) or receptor-like kinases (RLKs), akin to those in pathogen-associated molecular pattern (PAMP)-triggered immunity (PTI) (Wisser *et al*., 2005; Lacombe *et al*., 2005; Schweizer and Stein, 2011; Hurni *et al*., 2015; Schoonbeek *et al*., 2015; Nelson *et al*., 2018). PRRs like CHITIN ELICITOR RECEPTOR-LIKE KINASE 1 (CERK1), FLAGELLIN SENSING 2 (FLS2), and EF-TU RECEPTOR (EFR) detect PAMPs such as chitin, flg22, and elf18 respectively (Miya *et al*., 2007; Gómez-Gómez and Boller, 2000; Zipfel *et al*., 2006). Upon PAMP recognition, plants rapidly produce ROS like H_2_O_2_, followed by mitogen-activated protein kinase (MAPK) activation and defense gene induction. Measurement of ROS production by purified PAMPs has been used extensively in various laboratories to provide consistent and reliable PTI assessment and to study the relationship between PTI response and QDR (Vetter *et al*., 2012, Samira *et al*., 2020).

We aimed to identify broad-spectrum QDR loci in oilseed rape (*Brassica napus*) using a common panel of genotypes. *Brassica napus* is a major crop worldwide, producing edible oil, biodiesel, and animal feed protein. Disease susceptibility significantly impacts *B. napus* yields so enhanced resistance is a major breeding objective. Employing associative transcriptomics (AT) (Harper *et al*., 2012; Havlickova *et al*., 2018), we previously identified single nucleotide polymorphism (SNP) and gene expression marker (GEM) candidate loci linked to *B. napus* QDR against *Pyrenopeziza brassicae* (Fell et al., 2022). The GEM analysis utilizes transcriptomic data representing a baseline pre-infection expression level—a foundation for investigating constitutive QDR. The AT pipeline has been successfully applied to diverse phenotypes including pathogen resistance, flowering time, and seed glucosinolate content (Harper *et al*., 2012, Woodhouse *et al*., 2021, Dakouri *et al*., 2021; Roy *et al*., 2021). However, to our knowledge, AT has never been used to research the similarities and differences between the loci underlying resistance to multiple pathogens.

Three hypotheses underlie our study. Firstly, using a diverse *B. napus* panel— comprising a range of crop types including winter, semi-winter, and spring oilseed rape, swede, and kale, resulting in 219k SNP variants across the genome—we hypothesized there would be variable pathogen resistance within the population due to the high level of genetic and morphological variation. This variation might correlate with constitutive defenses or early innate immunity since PAMP recognition is conserved across pathogens. Therefore, our second hypothesis is that shared QDR loci may be associated with resistance to multiple pathogens, including those with contrasting lifestyles. Finally, given pathogens of different lifestyles utilize different mechanisms for their infection, our third hypothesis is that there would be some QDR loci dependent on the lifestyle of the pathogen. To test these hypotheses, we examined resistance to two necrotrophs, *Alternaria brassicicola* and *Botrytis cinerea,* and two hemibiotrophs, *P. brassicae,* and *Verticillium longisporum*, within the diversity panel. We used AT to identify the associated loci that define the “host genetic signature” of QDR in *B. napus*. Additionally, we explored the presence of loci associated with both QDR and PTI (measured by the production of PAMP-induced ROS).

For the first time, we demonstrate that AT analyses from several pathogens can be combined to identify candidate loci for broad-spectrum QDR. We identify GEMs associated with either resistance or susceptibility dependent on the pathogen lifestyle. Additionally, we identify a potential deletion associated with resistance to *V. longisporum* and potentially broad-spectrum QDR. This study provides new insight into the commonalities underlying broad-spectrum QDR and provides a resource for further mechanistic exploration and improving broad- spectrum resistance to pathogens in *B. napus*.

## Materials and Methods

### Pathogen infection assays

We selected a subset of the 191 *B. napus* genotypes for phenotyping, ensuring feasible pathology assays with ample replication across all participating labs. To mitigate environmental effects during seed production, all genotypes were cultivated under consistent environmental conditions before distribution to labs. Nevertheless, in some pathosystems, certain lines yielded unreliable data, leading to variable numbers of genotypes used for analyses.

#### Alternaria brassicicola

151 *B. napus* accessions were grown in peat soil (pH 5.6 – 6.8) with 1/30 Perlite, 16 h day/8 h night photoperiod under fluorescent light (Super TLD Philips 865, 100 μmol m^-2^ s^-1^), 19°C ± 2°C and approximately 70% relative humidity. The wild-type *A. brassicicola* strain (ATCC 96836) was cultured and previously described (Macioszek *et al*., 2018). A conidial suspension diluted to 3.5x10^5^ conidia ml^-1^ in distilled water was applied in two 10 μl drops to the third leaf of five-week-old plants. Necrotic areas were measured in three replicates per genotype at 5 days post-inoculation using WinDIAS_3 Imaging Analysis System (Delta-T Devices, UK). The experiment was repeated twice.

#### Botrytis cinerea

We divided 190 *B. napus* accessions into four batches for screening. Plants were grown in Levington F2 compost with 15% 4mm grit in growth chambers for 6-7 weeks under TL-tubes with a near-sunlight spectrum of 100 µmol m^-2^ s^-1^ with a photoperiod of 10 h. Temperatures were 20-22 °C/18-20 °C day/night. *Botrytis cinerea* strain B05.10 (Schoonbeek *et al*., 2001) was grown as described previously (Stefanato *et al*., 2009) and used for inoculations as described (Lloyd *et al*., 2014). Spore suspensions (200 µl at 2.5×10^6^ spores ml^-1^) were spread on 1/10 PDA (2.5 g l^-1^ potato dextrose broth, 12 g l^-1^ agar), and after 24 h at 21°C, 4 mm agar plugs were used for *B. napus* inoculations of 22 mm leaf discs on 0.6% water agar. Lesion diameter of 16 replicates per genotype was measured after 48 h at 21 °C, 85-100% relative humidity, and low light (10-20 µmol m^-2^ s^-1^). Two full experimental repeats were conducted.

#### Pyrenopeziza brassicae

The *P. brassicae* methods and phenotype dataset used were the same as in our previous publication (Fell *et al*., 2022). Disease severity was scored on a scale of 1 to 6—with a score of 1 for no sporulation and 6 for the most sporulation—and was calculated for the 129 genotypes.

#### Verticillium longisporum

We divided 191 *B. napus* accessions into four batches for screening. Six reference lines (Falcon, SEM, Zhongyou, Loras) were used for normalization. Each line had 20 plants for both mock and *V. longisporum* inoculation. The preparation of fungal inoculum and disease assessment was done as described previously (Zheng, 2018; Zheng *et al*., 2019). Ten-day-old conidial suspension of *V. longisporum* isolate VL43 obtained from a diseased *B. napus* plant (Zeise and Tiedemann, 2001) was used for inoculation. Ten-day-old plants with unfolded cotyledons were inoculated with water or 1x10^6^ CFU ml^-1^ spore suspension for 50 minutes. Plants were kept in a climate chamber with a 16-hour photoperiod and 22 ± 2°C temperature. Stem disease severity ranging from 1 (healthy) to 9 (dead) was assessed at 7, 14, 21, and 28 days post-inoculation. The area under the disease progress curve was calculated and normalized against reference lines.

### ROS assays

190 B. napus accessions were split into four groups and cultivated in Levington F2 compost with 15% 4 mm grit in growth chambers. A near-sunlight spectrum at around 100 µmol m-2 s-1 and a 10-hour photoperiod were maintained, with temperatures of 20-22°C/18-20°C day/night. ROS measurements employed a luminol/peroxidase-based assay. For each accession, two 4 mm diameter leaf discs were taken from the second most recently expanded leaf of four individual plants, incubated in 200 µl sterile water in a 96-well plate for 16-24 hours in darkness. Subsequently, water was removed and a solution was added containing 34 mg l-1 luminol, 20 mg l-1 horseradish peroxidase (HRP), and PAMP: either chitin (100 g l-1, NA-COS-Y, Yaizu Suisankagaku Industry CO, YSK, Yaizu, Japan); flg22 (QRLSTGSRINSAKDDAAGLQIA) or elf18 (SKEKFERTKPHVNVGTIG, both 10 mM, Peptron, http://www.peptron.co.kr).

Luminescence was recorded over 40 minutes, and the total ROS response was quantified using area under the curve calculations. Two replicate experiments were conducted.

### Phenotypic data transformation for association analysis

To maximize statistical power, all disease phenotype data per pathogen and *B. napus* cultivar were included (**Table S1**). Each dataset (infection phenotypes for *A. brassicicola, B. cinerea, P. brassicae,* or *V. longisporum,* ROS measurements for chitin, flg22, or elf18) was analyzed using a linear fixed- and random effects model with *B. napus* genotype as a fixed effect and experimental replicate and batch as random effects. Estimated marginal means were calculated for each *B. napus* genotype using emmeans (version 1.8.0) (R package) (R Core Team, 2023). These values were used for input into the AT pipeline. For combined data visualization, estimated marginal means were normalized to a 0-1 range. In pathogen datasets, the reciprocal of these values was used to represent ’resistance’.

### Associative transcriptomics

We used genotype (SNP) and expression level datasets (Havlickova *et al*., 2018), from York Knowledgebase (http://yorknowledgebase.info) and refined to include only the lines used within this study. Functional genotypes were determined via 100-base mRNAseq for the third true leaf using the Illumina HiSeq 2000 platform. Sequence reads were mapped to the CDS gene model-based Brassica AC pan-transcriptome reference (He *et al*., 2015), which comprised 116,098 gene models for SNP scoring and read quantification. 219,454 SNPs with maternal allele frequencies greater than 5% were used for downstream analysis. Genome-Wide Association (GWA) and GEM mapping were done using the R-based GWA and GEM Automation (GAGA) pipeline (Nichols, 2022), which utilizes GAPIT Version 3 (Lipka *et al*., 2012; Wang and Zhang, 2021). GAGA was run using our recently updated population structure (Fell *et al*., 2022) and *B. napus* Pantranscriptome version 11 (Havlickova *et al*., 2018). For GWA analyses, generalized linear model (GLM), Bayesian-information and Linkage-disequilibrium Iteratively Nested Keyway (BLINK) (Huang *et al*., 2019), and Fixed and random model Circulating Probability Unification (FarmCPU) (Liu *et al*., 2016) models were compared using QQ plots to select the best-fitting model.

GEM association analyses were performed within the GAGA pipeline based on methodology by Harper *et al*. (2012). Associations were determined by linear regression using Reads Per Kilobase of the transcript, per Million mapped reads (RPKM) to predict a quantitative outcome of the trait value. Markers with an average expression below 0.5 RPKM were excluded prior to analysis resulting in 53,883 expression values for association analysis. The Pearson coefficient was utilized to assess the correlation between expression and resistance phenotype for each GEM.

To determine the statistical significance threshold in GWA analysis, various methods that account for multiple testing have been proposed, including the Bonferroni correction, Sidak correction, and False Discovery Rate (FDR). Due to the prevalence of linked markers in modern GWA, the Bonferroni correction threshold can often have an inflated significance level, detecting only strong, major gene effects as significant. To address this, we utilized the FDR to manage the expected proportion of false positives among significant associations. FDR values for both GEM and GWA were determined using the q-value R package (Storey, 2011). Bonferroni thresholds for GWA and GEM are 6.64 and 6.03- Log_10_P respectively (P<0.05) and 6.34 and 5.73 -Log_10_P (P<0.1).

Linkage disequilibrium (LD) varies based on chromosome position and selection level. To ascertain the LD at the A09 locus associated with *V. longisporum* resistance, we calculated the mean pairwise R^2^ between this marker and all others on the chromosome using TASSEL Version 5.0 site by all analysis option (Bradbury *et al*., 2007). Markers were deemed in LD when *R^2^* > 0.2.

### Interpretation of data post-AT

We identified potential *Arabidopsis thaliana* orthologs of *B. napus* genes using BLASTN analysis against the *A. thaliana* transcriptome (TAIR version 10). Hypergeometric probability was used for to determine if the number of overlapping GEMs observed was greater than that expected by chance as detailed by Kim *et al*. (2001).

We conducted weighted gene co-expression analysis (WGCNA) using the R- based WGCNA library (v 1.72) (Langfelder and Horvath, 2008). Low expressed genes with 0 RPKM value in half of the samples were removed and rest of the data was used to perform a signed WGCNA analysis to detect modules of genes using the blockwiseModules() function. A soft threshold power of 6 was selected based on minimum mean connectivity using the pickSoftThreshold() function within the WGCNA package. Correlation and p-values with phenotypic trait data were calculated using the “Cor()” and “CorPvalueStudent()” functions. Gene Ontology (GO) enrichment analysis (FDR < 0.05) was conducted using a Fisheŕs exact test for GEMs within WCGNA modules.

## Results

### QDR to A. brassicicola, B. cinerea, P. brassicae and V. longisporum is present in the B. napus diversity panel

Given the extensive genetic diversity in our *B. napus* panel we hypothesized that QDR to *A. brassicicola, B. cinerea, P. brassicae,* and *V. longisporum* would be present. We did pathogenicity assays and found that the *B. napus* genotypes demonstrated varying levels of resistance to the pathogens, confirming the presence of QDR in the panel (**Fig. 1a**, **Table S1**).

**Fig. 1.**
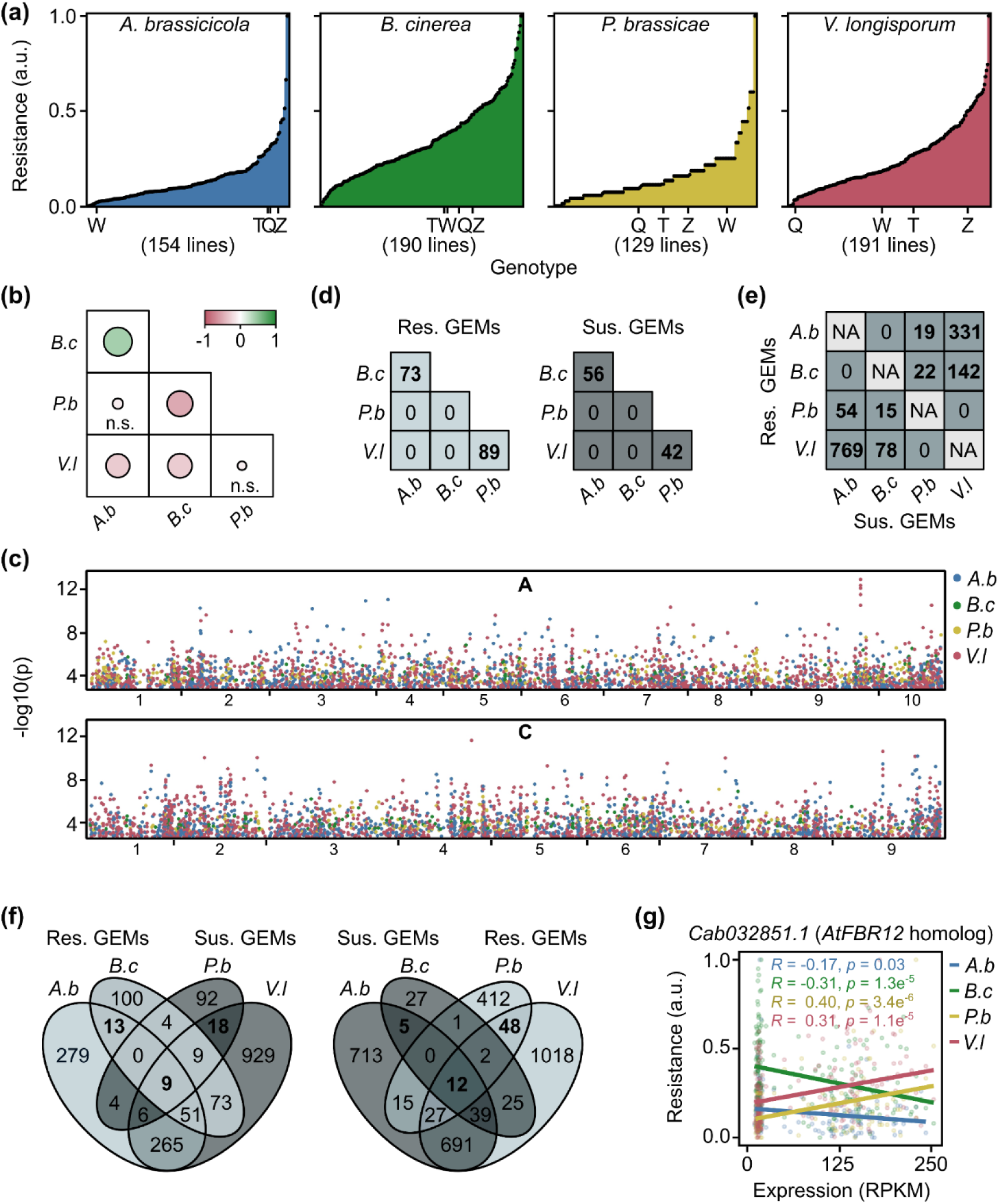
*Brassica napus* genotypes have common GEMs associated with resistance pathogens of the same lifestyle. **(a)** *B. napus* disease resistance to fungal pathogens is quantitative. Resistance phenotype (arbitrary units (a.u.), normalized values between zero and one) of different *B. napus* genotypes to *Alternaria brassicicola* (*A.b*), *Botrytis cinerea* (*Bc*), *Pyrenopeziza brassicae* (*Pb*) or *Verticillium longisporum* (*Vl*). The total number of genotypes used for each pathogen assay and the position of reference genotypes Quinta (Q), Tapidor (T), Westar (W) and Zhongshuang 11 (Z) are indicated. (**b**) Correlation between the resistance phenotype of *B. napus* lines to the fungal pathogens. Positive correlations (green), negative correlations (red) and no correlation (n.s.); strengths of correlation (sizes of circles) are indicated between pair-wise comparisons of resistance responses. (**c**) Manhattan plot of *B/ napus* genome showing marker-trait association of statistically significant GEMs for resistance to each fungal pathogen. The x-axis indicates GEM location along the chromosome; the y-axis indicates the -log_10_(p) (*P* value). (**d**) The numbers of resistance and susceptibility gene expression markers (GEMs) shared between pair-wise comparisons of pathogens. (**e**) The numbers of GEMs associated with resistance to one pathogen and susceptibility to another pathogen. (**f**) Venn diagrams showing the overlap between *A.b and B.c* susceptibility GEMs and *P.b* and *V.l* resistance GEMs (right) and the overlap between *A.b and B.c* resistance GEMs and *P.b* and *V.l* susceptibility GEMs (left). (**g**) Linear regression analysis of gene expression (RPKM) of *Cab032851.1* relative to resistance to fungal pathogens (arbitrary units (a.u.), normalized values between zero and one).

We hypothesized that if broad-spectrum QDR was present, there would be a positive correlation between phenotypic responses to pathogens. We initially examined disease resistance in four *B. napus* genotypes with published reference genomes: Quinta, Tapidor, Westar, and Zhongshuang 11. Quinta showed low resistance to both hemibiotrophic pathogens, while Zhongshuang 11 displayed high resistance to both necrotrophic pathogens (**Fig. 1a**). However, the ranking of these reference genotypes within the diversity panel varied. To explore correlations across the entire *B. napus* panel’s responses to pathogens, we generated a correlation matrix for pairwise fungal pathogen comparisons (**Fig. 1b**). There was a positive correlation between resistance to necrotrophic pathogens *A. brassicicola* and *B. cinerea*. Conversely, negative correlations emerged between resistance to different pathogen lifestyles; for example, hemibiotrophic *V. longisporum* showed negative correlations with necrotrophic pathogens *A. brassicicola* and *B. cinerea*.

### GEMs are associated with resistance to pathogens of the same lifestyle

To identify QDR loci, we performed GWA analysis on our datasets from the four fungal pathogens. Minor GWA association peaks could be observed (-Log_10_P>5) however we did not identify any significant marker associations with resistance to any of the pathogens, using either a Bonferroni threshold of P=0.1 or a false discovery rate (FDR) of 0.05. All association data provided is within **Table S2**. We performed GEM analyses using the transcriptomes of the *B. napus* genotypes to associate resistance with the expression of all gene models (**Table S3**). Numerous GEMs showed significant associations with fungal pathogen resistance (FDR < 0.05), and these were distributed evenly across the *B. napus* genome (**Fig. 1c**): 2129 GEMs for *A. brassicicola*, 370 for *B. cinerea*, 659 for *P. brassicae*, and 3222 for *V. longisporum*. We classified GEMs based on their positive or negative correlation with resistance or susceptibility, naming them ’resistance GEMs’ or ’susceptibility GEMs,’ respectively (**Table 1**).

**Table 1.** The number of gene expression markers (GEMs) associated with resistance to *Brassica napus* fungal pathogens *Alternaria brassicicola, Botrytis cinerea, Pyrenopeziza brassicae and Verticillium longisporum.* The total numbers of GEMs, GEMs with expression positively correlated with resistance (R, resistance) and negatively correlated with resistance (S, susceptibility) are indicated.

| Pathogen | Number of GEMs (FDR < 0.05) |  |  |
| --- | --- | --- | --- |
|  | Total | R | S |
| *A. brassicicola* | 2129 | 627 | 1502 |
| *B. cinerea* | 370 | 259 | 111 |
| *P. brassicae* | 659 | 517 | 142 |
| *V. longisporum* | 3222 | 1862 | 1360 |

We hypothesized that there might be shared QDR GEMs associated with all four fungal pathogens. To investigate this, we compared the lists of resistance and susceptibility GEMs across the fungal pathogens (**Fig. 1d**). While necrotrophic pathogens (*A. brassicicola* and *B. cinerea*) had shared GEMs, as did hemibiotrophic pathogens (*P. brassicae* and V. *longisporum*), there were no common GEMs between necrotrophic and hemibiotrophic pathogens. Consequently, there were no shared GEMs associated with broad-spectrum resistance spanning pathogens with differing lifestyles.

Given the existence of shared QDR GEMs associated with pathogens of the same lifestyle and that pathogens with similar lifestyles have similarities related to their infection, feeding strategies, and plant processes required for defense, we conducted overrepresentation analyses to compare the number of shared QDR GEMs between pathogens of the same lifestyle against what would be expected by chance (Plaisier *et al*., 2010). There was significant overrepresentation in the number of shared GEMs between necrotrophic pathogens *A. brassicicola* and *B. cinerea* (73 resistance GEMs and 56 susceptibility GEMs), as well as between hemibiotrophic pathogens *P. brassicae* and *V. longisporum* (89 resistance GEMs and 42 susceptibility GEMs) (**Fig. 1d, Table S4, S5**).

### GEMs are associated antagonistically with resistance to pathogens of different lifestyles

Hemibiotrophic and necrotrophic pathogens employ distinct plant mechanisms for their infection, with certain genes that confer resistance to hemibiotrophic pathogens leading to susceptibility against necrotrophic pathogens (Lorang *et al*., 2007). Thus, we hypothesized the presence of QDR GEMs with antagonistic associations based on pathogen lifestyle. To explore this, we compared resistance GEMs with susceptibility GEMs for each combination of fungal pathogens (**Fig. 1e**). As anticipated, we identified no GEMs with antagonistic effects for pairs of either necrotrophic or hemibiotrophic pathogens. However, intriguingly, we observed an overrepresentation of GEMs with antagonistic resistance associations between necrotrophic and hemibiotrophic pathogens. For instance, 1100 (769 + 331) GEMs with opposite resistance associations were linked to *A. brassicicola* and *V. longisporum* (**Table S4, S5**).

Broad-spectrum resistance GEMs with no antagonistic effects on susceptibility to other pathogens hold promise for crop breeding. To identify such candidates, we compared necrotrophic pathogen susceptibility GEMs to hemibiotrophic pathogen resistance GEMs. Similarly, we compared necrotrophic pathogen resistance GEMs to hemibiotrophic pathogen susceptibility GEMs (**Fig. 1f, Table S4**). 13 GEMs were associated with resistance to both *A. brassicicola* and *B. cinerea* with no associations with susceptibility to *P. brassicae* and *V. longisporum* (**Table 2**). 48 GEMs were associated with resistance to both *P. brassicae* and *V. longisporum* with no associations with susceptibility to *A. brassicicola* and *B. cinerea* (**Table 3**). Notably, four GEMs homologous to *A. thaliana ACIP1* were associated with resistance to both hemibiotrophic pathogens and no antagonistic associations with susceptibility to the necrotrophic pathogens.

**Table 2.** Broad-spectrum GEMs associated with resistance (R) or susceptibility (S) to necrotrophic fungi (necro.) *Alternaria brassicicola* and *Botrytis cinerea,* with no antagonistic effects on hemibiotrophic fungi (hemi.) *Pyrenopeziza brassicae* and *Verticillium longisporum.* The putative *Arabidopsis thaliana* (*At*) orthologs, protein class, and gene ID are indicated.

| GEM | Necro. | Hemi. | At ortholog | Gene name | Protein class |
| --- | --- | --- | --- | --- | --- |
| Cab019641.1 | R | - | - | - | - |
| Cab029666.1 | R | - | AT1G04980.1 | *PDI10; Protein disulfide-isomerase like 2-2* | chaperone |
| Cab006731.1 | R | - | - | - | - |
| Cab023848.3 | R | - | AT3G13772.1 | *TMN7; Transmembrane 9 superfamily member 7* | transporter |
| Bo6rg071140.1 | R | - | AT3G56460.1 | *GroES-like zinc-binding alcohol dehydrogenase family protein* | oxidoreductase |
| Cab006719.1 | R | - | AT5G24630.5 | *BIN4; Brassinosteroid insensitive 4* | DNA-binding |
| Bo3g028280.1 | R | - | AT4G38220.2 | *AQI, Aquaporin interactor, N-acyl-L-amino-acid amidohydrolase* | - |
| Bo9g183340.1 | R | - | AT5G02820.1 | *BIN5, Brassinosteroid insensitive 5, DNA topoisomerase 6 subunit A* | endodeoxyribonuclease |
| Bo3g004890.1 | R | - | AT5G08380.1 | *AGAL1; Alpha-galactosidase 1* | galactosidase |
| Cab010087.2 | R | - | - | - | - |
| Cab023861.2 | R | - | AT1G72730.1 | *Eukaryotic initiation factor 4A-3* | RNA helicase |
| BnaC03g41840D | R | - | - | - | - |
| Bo1g017790.1 | R | - | AT4G17720.1 | *BPL1; Putative RRM-containing protein* | RNA splicing factor |
| Cab007046.1 | S | - | AT5G48590.1 | - | - |
| Bo5g132170.1 | S | - | AT3G13740.1 | *RNC4; RNase III-like enzyme* | - |
| Cab029917.1 | S | - | AT1G01090.1 | *PDH-E1 ALPHA; Pyruvate dehydrogenase E1 subunit alpha-3* | dehydrogenase |
| Bo3g040470.1 | S | - | AT2G27510.1 | *FD3; Ferredoxin-3* | reductase |
| Bo5g132390.1 | S | - | AT3G13550.1 | *COP10, Constitutive photomorphogenesis protein 10* | ubiquitin-protein ligase |

**Table 3.** Broad-spectrum GEMs associated with resistance (R) or susceptibility (S) to hemibiotrophic fungi (hemi.) *Pyrenopeziza brassicae* and *Verticillium longisporum,* with no antagonistic effects on necrotrophic fungi (necro.) *Alternaria brassicicola* and *Botrytis cinerea.* The putative *Arabidopsis thaliana* (*At*) orthologs, protein class, and gene ID are indicated. Multiple GEMs corresponding to *A. thaliana* ortholog AT3G09980.1 are highlighted.

| GEM | Necro. | Hemi. | At ortholog | Gene name | Protein class |
| --- | --- | --- | --- | --- | --- |
| BnaC09g43580D | - | R | - | - | - |
| Cab025354.2 | - | R | AT2G45580.1 | CYP76C3; Cytochrome P450, family 76, subfamily C, polypeptide 3 | - |
| Bo4g182630.1 | - | R | AT2G34470.1 | UREG; Urease accessory protein G | - |
| **Bo5g058580.1** | - | **R** | **AT3G09980.1** | **ACIP1; Acetylated interacting protein 1** | - |
| Bo4g183100.1 | - | R | AT2G35190.1 | NPSN11; Novel plant SNARE 11 | SNARE protein |
| Bo2g095510.1 | - | R | AT1G80930.1 | Pre-mRNA splicing factor CWC22 homolog | RNA processing factor |
| Bo9g176470.1 | - | R | AT5G06250.2 | NGAL3; NGATHA-like protein 3, B3 domain-containing protein | transcription factor |
| **BnaC03g64760D** | - | **R** | **AT3G09980.1** | **ACIP1; Acetylated interacting protein 1** | - |
| Cab042580.1 | - | R | AT4G02710.1 | NET1C; Protein NETWORKED 1C, Ki-se interacting (KIP1-like) | - |
| Bo4g045470.1 | - | R | AT2G32860.1 | BGLU33; beta glucosidase 33 | - |
| Bo4g182440.1 | - | R | AT2G34250.2 | SecY/Sec61-alpha family member protein | transporter |
| Cab042707.1 | - | R | AT5G51120.1 | PAMN1; Polyadenylate-binding protein 1 homolog | translation initiation factor |
| Cab037277.2 | - | R | AT3G22550.1 | FLZ8; FCS-Like Zinc finger 8 | - |
| Bo4g183310.1 | - | R | AT2G35510.1 | SRO1, Similar to RCD 1, I-ctive poly [ADP-ribose] polymerase | - |
| Cab025625.4 | - | R | - | - | - |
| Bo6rg103690.1 | - | R | AT1G67280.2 | GLYI6; Lactoylglutathione lyase | lyase |
| Bo4g182680.1 | - | R | - | - | - |
| Bo2g095560.1 | - | R | - | - | - |
| Cab029121.1 | - | R | AT2G26250.1 | KCS10; 3-ketoacyl-CoA synthase 10 | - |
| **Bo5g137820.1** | - | **R** | **AT3G09980.1** | **ACIP1; Acetylated interacting protein 1** | - |
| Bo4g194830.1 | - | R | AT2G44120.1 | 60S ribosomal protein L7-3 | ribosomal protein |
| Cab013800.1 | - | R | AT1G16445.1 | S-adenosyl-L-methionine-dependent methyltransferases superfamily | methyltransferase |
| Bo2g095550.1 | - | R | - | - | - |
| Bo4g182550.1 | - | R | AT2G34410.2 | RWA3; Reduced wall acetylation 3 homolog | - |
| Bo4g182960.1 | - | R | AT2G34840.1 | epsilon2-COP; Coatomer subunit epsilon-2 | vesicle coat protein |
| Bo1g051850.1 | - | R | AT2G31290.2 | Ubiquitin carboxyl-terminal hydrolase family protein | - |
| BnaC06g39270D | - | R | - | - | - |
| Bo1g003050.1 | - | R | AT4G40000.1 | TRM4A; S-adenosyl-L-methionine-dependent methyltransferase | R- methyltransferase |
| Bo4g023900.1 | - | R | AT2G44620.1 | Acyl carrier protein 1, mitochondrial | - |
| Cab002494.2 | - | R | AT2G30320.1 | - | - |
| Cab007584.1 | - | R | AT5G05800.2 | L10-interacting MYB domain-containing protein | - |
| **BnaC04g02850D** | - | **R** | **AT3G09980.1** | **ACIP1; Acetylated interacting protein 1** | - |
| BnaC05g07480D | - | R | - | - | - |
| Bo5g003800.1 | - | R | AT1G03365.1 | Putative E3 ubiquitin-protein ligase RF4 | - |
| Cab042031.1 | - | R | - | - | - |
| Bo4g183270.1 | - | R | - | - | - |
| Bo4g183070.1 | - | R | AT2G35050.1 | *Kinase superfamily domain-containing protein* | non-receptor ser/thr kinase |
| Bo4g182310.1 | - | R | AT2G33860.1 | *ARF3/ETT; Auxin response factor 3/ETTIN* | - |
| Cab019919.4 | - | R | AT4G31040.1 | *DLDG1; Chloroplast envelope membrane protein* | - |
| Bo6rg003510.1 | - | R | AT3G18524.1 | *MSH2; DNA mismatch repair protein* | D- metabolism protein |
| Cab019786.1 | - | R | AT4G29530.1 | *THMPASE1; Thiamine phosphate phosphatase-like protein* | phosphatase |
| Bo4g182420.1 | - | R | AT3G18430.2 | *Calcium-binding EF-hand family protein* | - |
| Bo6rg003540.1 | - | R | AT1G44790.1 | *Gamma-glutamylcyclotransferase 2-3* | - |
| Cab007776.1 | - | R | AT5G11710.1 | *EPS1; Clathrin interactor EPSIN 1* | membrane trafficking |
| Bo4g183130.1 | - | R | AT2G35230.1 | *IKU1; Protein HAIKU1* | - |
| Bo4g143960.1 | - | R | AT5G37890.1 | *SI-L7; E3 ubiquitin-protein ligase SI--like 7* | ubiquitin-protein ligase |
| Cab007778.1 | - | R | - | - | - |
| Bo6rg003520.1 | - | R | AT1G44770.1 | *elongation factor* | - |
| Cab019945.1 | - | S | - | - | - |
| Cab044891.1 | - | S | - | - | - |
| BnaA08g28590D | - | S | - | - | - |
| Bo6rg033210.1 | - | S | AT1G55090.1 | *Glutamine-dependent -D(+) synthetase* | ligase |
| BnaC08g18360D | - | S | - | - | - |
| Bo2g007540.1 | - | S | AT5G07210.1 | *ARR21; Putative two-component response regulator 21* | transcription factor |
| BnaA01g02330D | - | S | - | - | - |
| Cab002672.1 | - | S | AT2G33810.1 | *SPL3; Squamosa promotor binding protein-like 3* | DNA-binding |
| Cab013528.1 | - | S | - | - | - |
| Cab034921.1 | - | S | AT2G38540.1 | *LP1; Non-specific lipid-transfer protein 1* | - |
| Cab020282.2 | - | S | - | - | - |
| Bo7g095690.1 | - | S | AT5G25830.1 | *GATA12; GATA transcription factor 12* | transcription factor |
| BnaC08g15170D | - | S | - | - | - |
| Cab037036.1 | - | S | - | - | - |
| Cab034089.1 | - | S | - | - | - |
| Bo8g087140.1 | - | S | AT3G54890.1 | *LHCA1; Light-harvesting complex gene 1* | - |
| Bo6rg098720.1 | - | S | AT1G67700.1 | *HHL1; Hypersensitive to high light 1* | - |
| Bo5g152840.1 | - | S | AT3G02150.2 | *PTF1; Plastid transcription factor 1* | transcription factor |

Interestingly, there was a significant overrepresentation of GEMs associated with susceptibility or resistance to both necrotrophic pathogens, but with opposing associations to both hemibiotrophic pathogens: 21 GEMs (9 + 12) (**Table 4**, S6). Notably, the expression of *Cab032821.1*, the homolog of *A. thaliana Fumosin B1- resistant 12, FBR12*, was associated with resistance to both hemibiotrophic pathogens but susceptibility to both necrotrophic pathogens (**Fig. 1g**), as was the *Bo8g081460.1,* the homolog of an *A. thaliana disease resistance nucleotide- binding leucine-rich repeat (NBS-LRR)* protein (**Table 4**).

**Table 4.** Broad-spectrum GEMs associated with resistance (R) or susceptibility (S) to hemibiotrophic fungi (hemi.) *Pyrenopeziza brassicae* and *Verticillium ongisporum,* with antagonistic effects on necrotrophic fungi (necro.) *Alternaria brassicicola* and *Botrytis cinerea.* The putative *Arabidopsis thaliana* (*At*) orthologs, protein class, and gene ID are indicated.

| GEM | Necro. | Hemi. | At ortholog | Gene name | Protein class |
| --- | --- | --- | --- | --- | --- |
| BnaA10g08980D | R | S | AT2G34470.2 | *UREG; Urease accessory protein G* |  |
| Bo2g101920.1 | R | S | AT3G10020.1 | *HUP26; Hypoxia unknown protein 26* |  |
| Cab002732.1 | R | S | - | - |  |
| Bo2g101910.1 | R | S | - | - |  |
| Bo4g021310.1 | R | S | AT2G44150.1 | *ASHH3; Histone-lysine N-methyltransferase* |  |
| BnaA06g22180D | R | S | - | - |  |
| Bo7g077710.1 | R | S | AT5G47570.1 | *NADH dehydrogenase ubiquinone 1 beta subcomplex subunit* | dehydrogenase |
| Cab031167.1 | R | S | - | - |  |
| Cab017493.1 | R | S | AT5G56940.1 | *Ribosomal protein S16 family protein* | ribosomal protein |
| Cab007887.1 | S | R | AT5G13310.1 | - |  |
| Bo6rg081580.1 | S | R | AT1G78420.2 | *DA2; mono-ubiquitinitor* |  |
| Bo9g176310.1 | S | R | AT5G06310.1 | *POT1b, Protection of telomeres 1b* | DNA metabolism protein |
| Cab029870.1 | S | R | AT1G02560.1 | *CLPP5; Nuclear encoded Clp protease 5* | serine protease |
| Bo6rg081770.1 | S | R | AT1G78170.1 | *EAU1, Ethylene-responsive element binding factor-associated* |  |
| Cab032816.1 | S | R | AT1G24170.1 | *LGT9, Galacturonosyltransferase-like 8* | glycosyltransferase |
| Cab027999.1 | S | R | - | - |  |
| Cab018896.1 | S | R | AT1G76610.1 | *MIZU-KUSSEI-like protein* |  |
| Bo4g151450.1 | S | R | - | - |  |
| Bo8g081460.1 | S | R | AT3G51570.1 | *Disease resistance protein (TIR-NBS-LRR class)* |  |
| Cab032851.1 | S | R | AT1G26630.1 | *FBR12; Fumonisin B1-resistant 12* | translation initiation factor |
| Bo4g149580.1 | S | R | AT2G22400.1 | *TRM4B, S-adenosyl-L-methionine-dependent methyltransferases* | RNA methyltransferase |

We conducted WGCNA analysis to explore the co-expression patterns of GEMs in our study. Several GEMs exhibited co-expression patterns across the *B. napus* population and were organized into co-expression modules. These modules were then assessed for their association with resistance to each pathogen (**Table S6**). Notably, significantly correlated modules included 22 modules (containing a total of 1075 GEMs) for resistance to *A. brassicola*, 18 modules (containing 299 GEMs) for resistance, 34 modules (containing 575 GEMs) for *P. brassicae* resistance, and 26 modules (containing 2075 GEMs) for *V. longisporum* resistance. Interestingly, modules correlated with resistance to necrotrophic pathogens displayed susceptibility to hemibiotrophic pathogens, and vice versa, consistent with the findings from our individual GEM results.

To gain further insights, we performed GO enrichment analysis on the putative *A. thaliana* orthologs of resistance and susceptibility GEMs within each module. Particularly noteworthy was a module linked to susceptibility to necrotrophic pathogens and resistance to hemibiotrophic pathogens, enriched in genes associated with photosynthesis. This suggests that photosynthesis-related processes might exert contrasting effects on resistance based on the pathogen’s lifestyle. We examined if there were any GEMs related to known pathogen-dependent pathways, specifically SA or JA signaling. We first used gene ontology to search for SA- or JA-related GEMs in our whole expression dataset (53,883 markers). No GEMs were linked to SA signaling. Markers associated with JA signaling were not enriched in our GEM analysis or WGCNA modules. This is likely because these genes are usually activated in response to pathogens rather than being constitutively expressed at the represented three-leaf stage in our expression data.

### GEMs associated with both PTI and QDR are dependent on pathogen lifestyle

We had originally hypothesized the presence of shared QDR loci associated with resistance to multiple pathogens with contrasting lifestyles possibly linked to early innate immunity due to the conserved nature of PAMP recognition. However, our comparisons of GEMs (**Fig. 1d**), revealed no shared GEMs between all four pathogens. Nevertheless, previous studies have linked PTI genes to QDR (Wisser *et al*., 2005; Schweizer and Stein, 2011; Hurni *et al*., 2015; Nelson *et al*., 2018). Thus, we proceeded to investigate whether any of our pathogen resistance QDR loci were related to PTI.

The *B. napus* genotypes demonstrated varying levels of PAMP-induced ROS production with all three PAMPs, suggesting quantitative traits (**Fig. 2a, S1a**). Reference lines (Quinta, Tapidor, Westar, and Zhongshuang 11) had comparable rankings for flg22 and elf18 ROS responses within the panel and our correlation matrix indicated that the *B. napus* genotypes exhibited similar rankings of responses across all three PAMPs (**Fig. S2b**). Minor GWA association peaks could be observed (-Log_10_P>5), with a single marker (Cab031873.1:1209:C) significantly associated with flg22-induced ROS production at a Bonferroni threshold of P=0.1. No significantly associated markers were observed at an FDR of 0.05. All marker associations are detailed in **Table S2**. We identified 184, 286, and 3251 GEMs associated with chitin-, flg22-, and elf18-induced ROS responses, respectively (**Table S3**), however, no GEMs corresponded to PRRs. There was a significant overlap between GEMs associated with the different PAMP responses. For example, out of the 184 chitin-induced ROS GEMs, 31% were shared with the flg22 response, 54% with the elf18 response, and 21% with both flg22 and elf18 responses (**Table S7, Fig. S1c**). These data suggest that response to PAMPs has some general genetic control and some PAMP-dependent distinct elements.

**Fig. 2.**
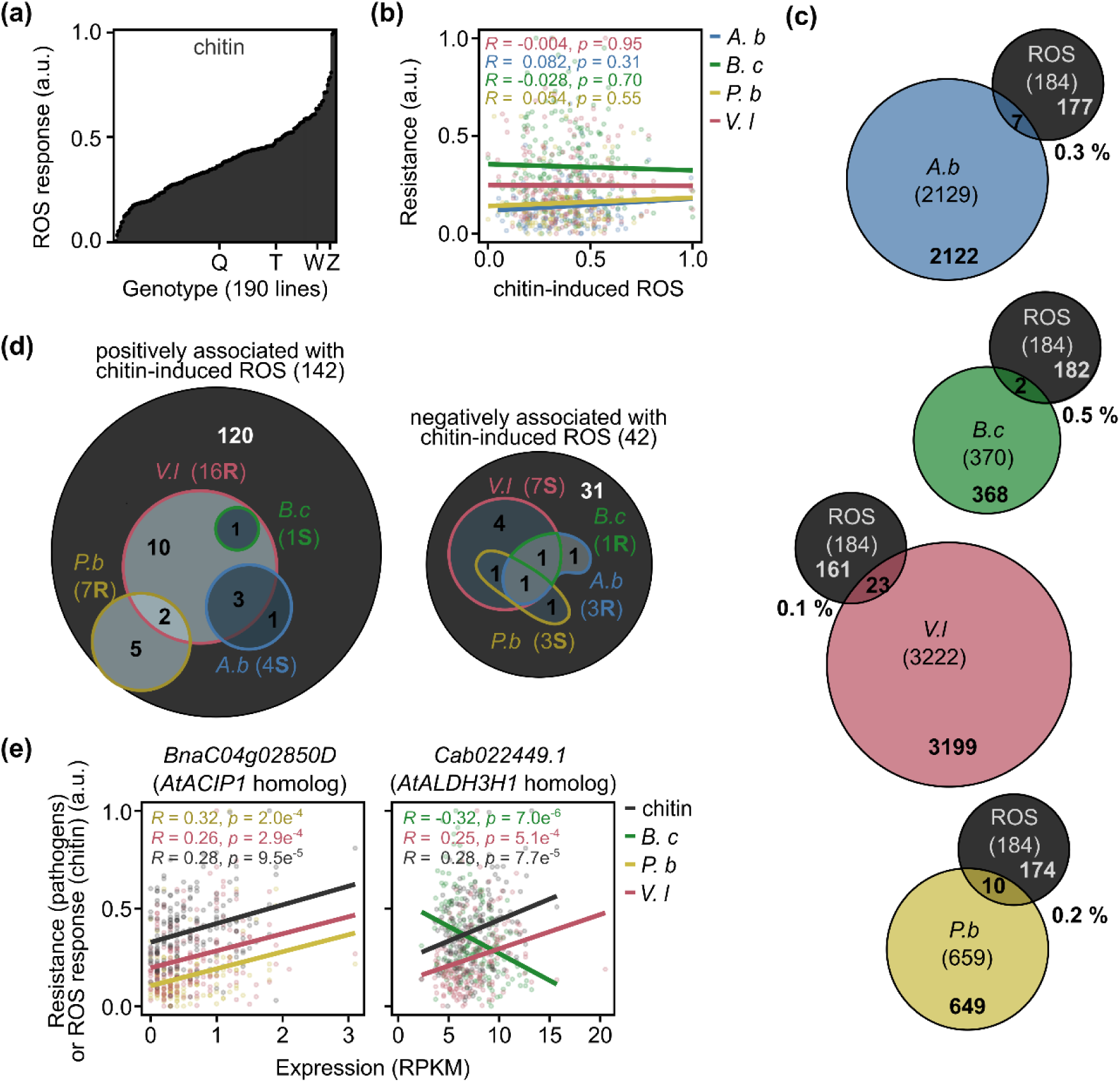
Shared GEMs for QDR and chitin-induced ROS are associated with resistance to hemibiotrophic pathogens but susceptibility to necrotrophic pathogens. (**a**) Chitin-induced ROS response (arbitrary units (a.u.), normalized values between zero and one) of different *B. napus* genotypes. The positions of reference genotypes Quinta (Q), Tapidor (T), Westar (W) and Zhongshuang 11 (Z) are indicated. (**b**) Linear regression analysis of chitin-induced ROS response relative to quantitative disease resistance to *Alternaria brassicicola (A.b), Botrytis cinerea (B.c)*, *Pyrenopeziza brassicae (P.b)* or *Verticillium longisporum (V.l)* (arbitrary units (a.u.), normalized values between zero and one). (**c**) Venn diagrams showing the overlap between GEMs associated with chitin-induced ROS and QDR to the fungal pathogens, *Alternaria brassicicola* (*A.b*), *Botrytis cinerea* (*B.c*), *Pyrenopeziza brassicae* (*P.b*), and *Verticillium longisporum* (*V.l*). The percentage of QDR-related GEMs which are also related to chitin-induced ROS is indicated. (**d**) The number of gene expression markers (GEMs) whose expression was positively or negatively associated with chitin-induced ROS and the number which were also associated with resistance (R) or susceptibility (S) to the *B. napus* fungal pathogens. Venn diagram indicating the overlap between GEMs associated with several pathogen interactions. (**e**) Linear regression analysis of gene expression (RPKM) of *BnaC04g2850D* and *Cab022449.1* relative to chitin-induced ROS responses and resistance to fungal pathogens.

Since chitin is a fungal PAMP and our study centres on fungal pathogens, we concentrated our following investigations on the overlap between chitin-associated and QDR-associated loci. Chitin-induced ROS response did not correlate with QDR to fungal pathogens (**Fig. 2b**) and only a small fraction (less than 1%) of QDR GEMs for each pathogen were linked to chitin-induced ROS response (**Table S7**). Notably, there was minimal overlap between GEMs associated with chitin-induced ROS response and QDR to necrotrophic pathogens, and a slight overlap with QDR to hemibiotrophic pathogens (**Table S5**). This suggests that most QDR loci are independent of early PAMP-induced ROS response (**Fig. 2c**).

Given that ROS accumulation can influence susceptibility to certain necrotrophic pathogens and our previous observation of QDR GEMs with lifestyle-dependent antagonistic associations, we investigated if loci linked to chitin-induced ROS production might be associated with resistance against hemibiotrophic pathogens but susceptibility to necrotrophic pathogens. Of the 184 GEMs that were associated with chitin-induced ROS, 142 were positively associated, i.e., increased gene expression was associated with the increased magnitude of chitin-induced ROS response. Remarkably, all chitin-induced ROS GEMs that coincided with QDR GEMs were exclusively associated with resistance to hemibiotrophic pathogens and susceptibility to necrotrophic pathogens (**Fig. 2d**, **Table 5**). These included *BnaC04g02850D*, a homolog of *A. thaliana ACIP1* (also associated with flg22 and elf18 responses), and *Cab022449.1*, a homolog of *A. thaliana Aldehyde dehydrogenase 3H1 (ALDH3H1)* (**Table S4**) (**Fig. 2e**). Likewise, among the 42 GEMs exhibiting negative correlations with chitin-induced ROS response, those overlapping with QDR were associated with resistance against necrotrophic pathogens and susceptibility to the hemibiotrophic pathogens. WGCNA analysis revealed that GEMs linked to chitin-induced ROS were grouped into 19 modules (comprising 135 GEMs) (**Table S6**). These modules, also showed associations with modules correlated with resistance to hemibiotrophic pathogens and susceptibility to necrotrophic pathogens, including the module enriched in GEMs related to photosynthesis regulation, aligning with our individual GEM results.

**Table 5.** The numbers of GEMs positively and negatively associated with chitin-induced ROS response and associated with either resistance (R) or susceptibility (S) to *Brassica napus* fungal pathogens *Alternaria brassicicola, Botrytis cinerea, Pyrenopeziza brassicae* or *Verticillium longisporum.* The direction of the correlation (positive or negative) of QDR GEMs compared to the chitin-induced ROS GEMs is also summarized.

| Pathogen | GEMs positively correlated with chitin-induced ROS (142) |  | GEMs negatively correlated with chitin-induced ROS (42) |  | Correlation to chitin-induced ROS GEMs |
| --- | --- | --- | --- | --- | --- |
|  | Number | Type (R/S) | Number | Type (R/S) |  |
| *A. brassicicola* | 4 | S | 3 | R | 7 -ve |
| *B. cinerea* | 1 | S | 1 | R | 2 -ve |
| *P. brassicae* | 7 | R | 3 | S | 10 +ve |
| *V. longisporum* | 16 | R | 7 | S | 23 +ve |

### A 0.51 MB deletion on chromosome A09 is associated with *V. longisporum* resistance and potentially broad-spectrum QDR

The genomic distribution of GEMs (**Fig. 1c**) highlighted a notable peak of five highly significant GEMs associated with *V. longisporum* resistance on chromosome A09 (*Cab013522.1, Cab013524.1, Cab013523.1, Cab013526.1,* and *Cab013517.1*). Notably, Cab013522.1, Cab013524.1, and Cab013526.1 were also linked to *P. brassicae* resistance, while Cab013522.1, Cab013523.1, Cab013524.1, and Cab013526.1 were associated with susceptibility to *A. brassicicola* (**Table S3**). This clustering suggests a potential region of interest for broad-spectrum QDR. Compellingly, this region of chromosome A09 also corresponded to a minor association (*P*=5.14e^-07^, FDR=0.11) from the *V. longisporum* resistance GWA analysis (**Fig. 3a**). Assessment of phenotypic variation segregating with alleles for Cab013526.1.135.A revealed that accessions inheriting a “G” at this locus had resistance values 57% lower than those inheriting an “A” (**Fig. 3b**). These alleles also segregated with resistance to *P. brassicae* (with “G” alleles showing 12% lower resistance than “A” alleles) and susceptibility to *A. brassicae* (with “G” alleles indicating 28% higher resistance than “A” alleles) (**Fig. 3b**). We calculated the distance in LD with this marker and found a region of 0.51 MB containing 107 genes (**Fig. 3c**).

**Fig. 3.**
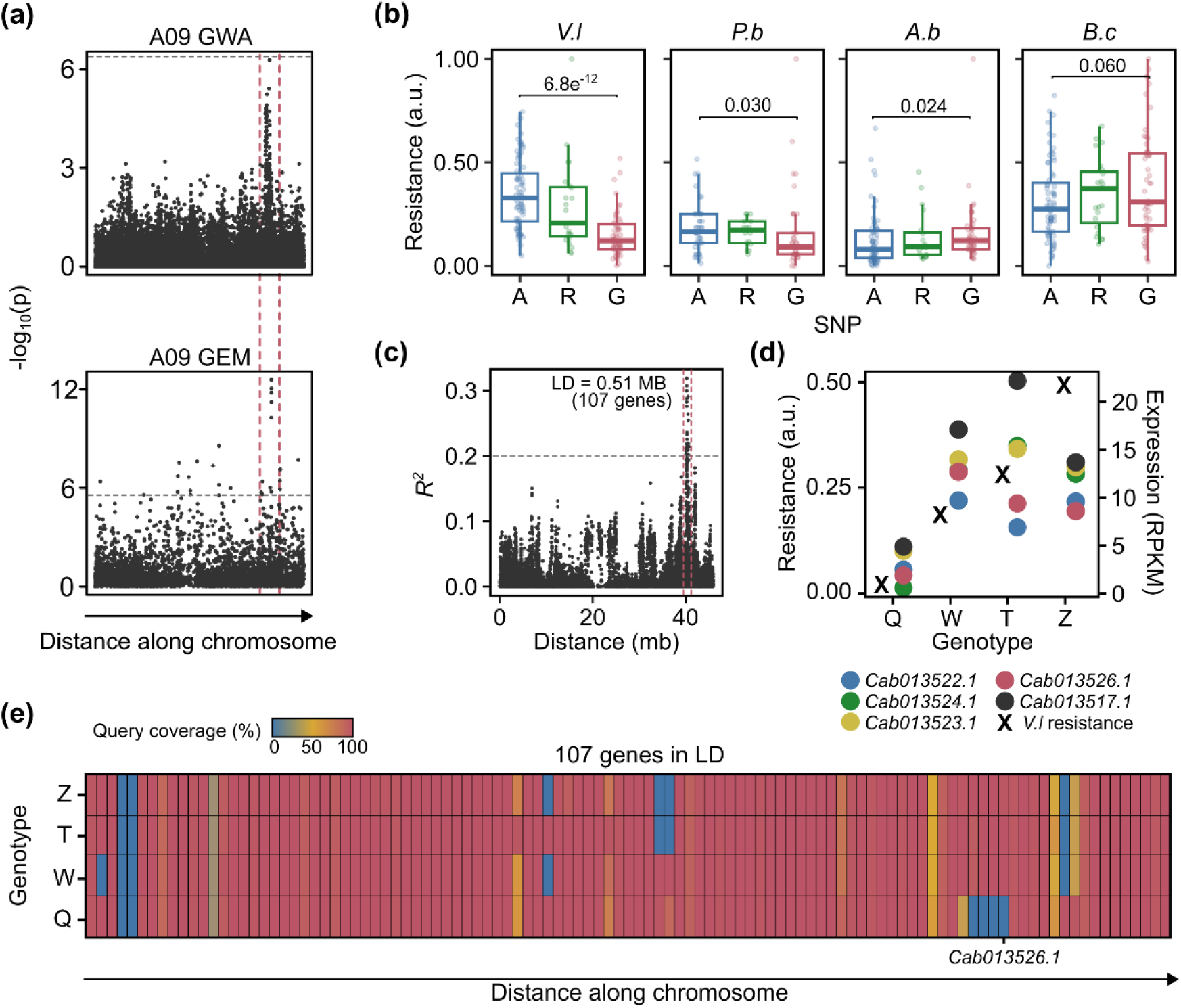
Resistance *to Verticillium longisporum* is associated with a genomic deletion on chromosome A09. (**a**) Manhattan plots showing marker-trait association resulting from GEM and GWA analysis of resistance to *V. longisporum* in 191 *Brassica napus* genotypes. The x-axis indicates GEM or SNP location along the chromosome; the y-axis indicates the -log_10_(p) (*P* value). Grey line indicates the FDR > 0.05 cut-off value, red line highlights the shared region. (**b**) Segregation of resistance to *V. longisporum (V.l), Pyrenopeziza brassicae (P.b), Alternaria brassicicola (A.b),* or *Botrytis cinerea (B.c)* (arbitrary units (a.u.), normalized values between zero and one) with the highest associating marker, Cab013526.1.651.A, showing the marker effect between A and G (R = A/G). *P* values were determined by a Student’s *t-*test. (**c**) Linkage decay plot from marker Cab013526.1.651.A as a function of genetic distance (MB). The grey line indicates an *R^2^* value of 0.2, red lines indicate the area of linkage disequilibrium. (**d**) Resistance to *V. longisporum* (left-hand y-axis) and gene expression (RPKM) of the five highest associating GEMs *Cab013522.1, Cab013524.1, Cab013523.1, Cab013526.1* and *Cab013517.1* (right-hand y-axis) in reference genotypes Quinta (Q), Tapidor (T), Westar (W) and Zhongshuang 11 (Z). (**e**) Heatmap indicating query coverage (compared to the *B. napus* pantranscriptome) in Q, T, W and Z for the 107 genes predicted to be in linkage disequilibrium with Cab013526.1.651.A on chromosome A09.

Given the association between the expression of five nearby GEMs and *V. longisporum* resistance, we speculated that this link could partly be due to a genomic deletion in this region. To explore this, we examined if any *B. napus* genotypes with published genomes (Quinta, Tapidor, Westar, and Zhongshuang 11 exhibited low gene expression and high susceptibility (Song *et al*., 2020). Quinta had little resistance to *V. longisporum* along with low expression of the five GEMs (expression at such levels could be a result of cross-mapping to the homoeologous region on the *B. napus* ‘C’ genome, C08) (**Fig. 3d**). On the other hand, Tapidor, Westar, and Zhongshuang 11 exhibited moderate resistance and gene expression levels. We hypothesized that Quinta has a deletion on *B. napus* chromosome A09, whereas Tapidor, Westar, and Zhongshuang 11 have not.

To determine the query coverage of the 0.51 MB region (107 genes) in LD with the most significant SNP marker, we conducted a BLAST analysis against the *B. napus* pantranscriptome for the 0.51 MB for each reference genotype. Notably, Tapidor, Westar, and Zhongshuang 11 showed nearly 100% query coverage for consecutive five genes—Cab013522.1 (40%), Cab013523.1 (0%), Cab013524.1 (0%), Cab013525.1 (0%), Cab013526.1 (0%)—while Quinta displayed reduced coverage (**Table S8**). BLAST analysis of the homoeologous region on chromosome C08 in Quinta revealed no differences compared to the other genotypes, suggesting a specific deletion on chromosome A09 (**Fig. S2a**). Linear regression analysis indicated that the expression of these five genes varied across *B. napus* genotypes and correlated with *V. longisporum* resistance (**Fig. S2b**), implying both quantitative and qualitative resistance traits associated with this region.

To identify potential genes within this deletion region that could contribute to V*. longisporum* resistance, we proposed that their homologs in the *B. napus* genome might also be relevant GEMs for resistance. Each of the five candidate genes on A09 only had one homolog, on chromosome C08. None of the five homologs were significant GEMs; however, linear regression analysis showed a significant association between gene expression of the *Cab013522.1* homolog (*Bo8g101810.1*) and the *Cab013524.1* homolog (Bo8g101830.1) and V. longisporum resistance (**Fig. S2b**). Downstream of the five genes, were two additional genes with nearly 100% sequence coverage in Quinta but reduced coverage in Tapidor, Westar, and Zhongshuang 11: *BnaA09g43290D* (0%) and *Cab013532.1* (40%). However, neither of these genes was expressed during the third-leaf stage used for GEM analysis. The corresponding C08 homolog of *Cab013532.1* (*Bo8g101910.1*) also was not expressed at this stage and no C08 homolog for *BnaA09g43290D* was identified.

To infer the potential function of the five gene candidates we identified their putative *A. thaliana* orthologs using BLAST (**Table 6**). The orthologs included a microtubule-binding motor protein, *Kinesin motor family protein, KIN7.2* (Cab013522.1); a carbohydrate kinase, *Xylulose kinase 1, XK1* (Cab013523.1); a multidrug and toxin extrusion (MATE) transporter, *Enhanced disease susceptibility 5 homolog, EDS5H* (Cab013524.1); a plastid isoform aldolase, *Fructose-bisphosphate aldolase 1, FBA1* (Cab013525.1); and a B-box zinc finger family transcription factor, *B-box domain protein 18, BBX18* (Cab013526.1).

**Table 6.** The five genes in the deletion on chromosome A09 associated with resistance against *Verticillium longisporum*. The putative *Arabidopsis thaliana* (*At*) orthologs, protein class, and gene ID are indicated.

| GEM | At ortholog | Gene name | Protein class |
| --- | --- | --- | --- |
| Cab013522.1 | AT2G21380.1 | **KIN7.2**; *Kinesin motor family protein* | microtubule binding motor protein |
| Cab013523.1 | AT2G21370.2 | **XK1**, *Xylulose kinase 1* | carbohydrate kinase |
| Cab013524.1 | AT2G21340.2 | **EDS5H**, *Enhanced disease susceptibility 5 homolog* | transporter (MATE efflux protein) |
| Cab013525.1 | AT2G21330.1 | **FBA1**; *Fructose-biphosphate aldolase 1* | aldolase |
| Cab013526.1 | AT2G21320.1 | **BBX18**; *B-box domain protein 18* | zinc family transcription factor |

## Discussion

### QDR is dependent on pathogen lifestyle

In this study, we used an AT pipeline to investigate constitutive QDR to *B. napus* fungal pathogens: *A. brassicicola, B. cinerea, P. brassicae,* and *V. longisporum* and define the “host genetic signature” of common and unique loci associated with resistance or susceptibility.

We initially hypothesized broad-spectrum QDR across multiple pathogens with contrasting lifestyles. However, our findings revealed shared QDR loci only within similar pathogen lifestyles, either hemibiotrophic or necrotrophic. This distinction likely stems from differing infection strategies. In fact, phenotypic resistance to necrotrophic pathogens was inversely related to resistance against hemibiotrophic pathogens. This contrast was mirrored by 21 GEMs associated with broad-spectrum resistance or susceptibility to necrotrophic pathogens with an opposite association with hemibiotrophic pathogens.

Immune responses, such as cell death, can effectively counteract the endophytic growth of some pathogens like *P. brassicae* and *V. longisporum,* but necrotrophic pathogens like *A. brassicicola* and *B. cinerea* may exploit cell death mechanisms to promote their infection (McCombe *et al*., 2022). Such contrasting effects have occasionally been previously observed, for example, an NBS-LRR gene conferring resistance to biotrophic pathogen *Puccinia coronata* f. sp. *avenae* acts as a susceptibility factor (*LOV1)* for necrotrophic pathogen *Cochliobolus victoriae* (Lorang *et al*., 2007). Compellingly, in our study, the ortholog of a novel *A. thaliana* NBS-LRR, *Bo8g081460.1*, was associated with resistance to both hemibiotrophic pathogens and susceptibility to both necrotrophic pathogens, as was the predicted *A. thaliana* ortholog of *FBR12* which was previously shown to be involved in cell death induced by *Pseudomonas syringae* (Hopkins *et al*., 2008).

We explored the overlap between QDR loci and PTI, and their varying associations with resistance based on pathogen lifestyle. While most QDR GEMs didn’t relate to PTI, those that were positively associated with chitin-induced ROS response were associated with resistance to hemibiotrophic fungi but susceptibility to necrotrophic fungi. For instance, the *B. napus* ortholog of A. thaliana ALDH3H1 positively correlated with chitin-induced ROS response and *V. longisporum* resistance, yet negatively correlated with resistance to *B. cinerea*. ALDH proteins—for example, a homolog of *A. thaliana* ALDH3H1, ALDH3I1— can mitigate oxidative stress by scavenging ROS (Kotchoni *et al*., 2006). Rapid ROS production during PTI can induce programmed cell death, which can confer susceptibility to necrotrophic pathogens like *B. cinerea* (Levine *et al*., 1994; Kotchoni and Gachomo, 2006). *B. cinerea* even produces ROS to promote its infection (Govrin and Levine, 2000) and ROS scavengers reduce infection severity (Tiedemann, 1997). Thus, it follows that genotypes with a potentially greater ability to scavenge ROS would be less susceptible to *B. cinerea*.

It’s intriguing that susceptibility to necrotrophic pathogens, resistance to hemibiotrophic pathogens, and ROS production aligned with a coexpressed module enriched in GEMs linked to photosynthesis regulation. Given that ROS are generated extensively during photosynthesis, it’s reasonable to suggest that some genes influencing photosynthesis could impact ROS production levels, consequently influencing resistance to necrotrophic and hemibiotrophic pathogens. Indeed, previous studies have noted distinct regulation of photosynthesis-related genes during *A. brassicicola* infection (Macioszek et al., 2020).

### Identification of a narrow chromosomal region associated with broad-spectrum QDR

By combining SNP and gene expression data through AT, we identified a novel locus associated with resistance to *V. longisporum*. Further in silico analyses of the region indicated five potential candidate genes, any of which could underly the resistance phenotype. Alleles of the BBX18 homolog also segregated with resistance to *P. brassicae* and KIN7.2, EDS5H, and BBX18 homologs were GEMs for resistance to *P. brassicae,* suggesting these could be associated with broad-spectrum QDR. Although we investigated this region using reference genotype with a genomic deletion (qualitative resistance), the expression of the KIN7.2, XK1, EDS5H, FBA1, and BBX18 homologs within other genotypes in the *B. napus* population varied and was significantly associated with resistance suggesting this locus is related to QDR.

KIN7.2 is involved in microtubule-based movement and—in the animal field— kinesins have been implicated in protein trafficking for anti-fungal defense (Kurowska *et al*., 2012; Ogbomo *et al*., 2018). EDS5H is a homolog of EDS5, a MATE protein which functions in plant salicylic acid (SA) dependent defense (Nawrath *et al*., 2002). However, unlike EDS5, in *A. thaliana* EDS5H is constitutively expressed in green tissues independent of pathogen infection and does not contribute to pathogen-induced SA accumulation (Parinthawong *et al*., 2015). Nevertheless, MATE proteins transport a broad range of, such as organic acids, plant hormones, and secondary metabolites, which could influence QDR (Nawrath *et al*., 2002; Takanashi, *et al*., 2014; Su, *et al*., 2022). Conversely, BBX18, as well as the additional XK1 and FBA1 genes in the potential deletion linked to resistance, lack documented defense associations.

With our in-silico analyses, we demonstrate the power of the AT pipeline to quickly narrow down on a predicted region of interest for broad-spectrum QDR. Additional functional analyses such as gene editing are needed to validate and identify the causal gene in *B. napus* and how it affects resistance to *V. longisporum* and other pathogens during infection.

### Broad-spectrum QDR is an opportunity for crop breeding

Achieving broad-spectrum disease resistance without compromising resistance to other pathogens is a crucial goal in crop breeding. For example, GEMs associated with susceptibility and no antagonistic associations with resistance could be interesting targets for gene editing to improve disease control without the introduction of transgenes (Hua *et al*., 2019). In addition, considering crop breeding for resistance, we identified GEMs associated with resistance to both necrotrophic pathogens and no antagonistic associations with susceptibility to the hemibiotrophic pathogens. We also identified GEMs associated with resistance to both hemibiotrophic pathogens and no antagonistic associations with susceptibility to the necrotrophic pathogens including four GEMs homologous to *A. thaliana ACIP1*— ACIP1 was previously implicated in flg22-induced ROS response and resistance to *Pseudomonas syringae* (Cheong *et al*., 2014). One of these four was positively associated with chitin-induced ROS response, highlighting it as a candidate for further investigation into its role in PTI and broad-spectrum QDR. Future work could extend our findings to evaluate how broad the spectrum of disease control is likely to be using specific fungal strains or mutants, other *B. napus* pathogens, and different crop species.

### Conclusions

Here, we used an AT pipeline to identify novel loci involved in QDR to several *B. napus* fungal pathogens *A. brassicicola, B. cinerea, P. brassicae* and *V. longisporum.* We observed broad-spectrum QDR loci were dependent on the lifestyle of the pathogen, as were associations with GEMs related to PAMP-induced ROS production. By combining GWA and GEM analyses we were able to identify a novel region of interest for *V. longisporum* resistance and potentially broad-spectrum QDR. To our knowledge, this is the first time this approach with several pathosystems has been used to identify loci involved in broad-spectrum QDR. In summary, this study provides new insight into broad-spectrum QDR and highlights interesting targets for crop breeding.

## Acknowledgments

The authors thank Paweł Jedyński and Tomasz Jęcz (University of Lodz, Poland) for help with plant cultivation and inoculation with *A. brassicicola* and Aiming Qi for advice about *P. brassicae* data analysis. VKM and AKK were supported by the National Center for Research and Development, Poland (grant no. ERA-CAPS II/1/2015). XZ and AvT were supported by the German Research Foundation (grant no. ERA-CAPS DFG; TI170/13–1). HJS, RW, RB, CJR, HF, GKM, HUS and BDLF were funded by the Biotechnology and Biological Sciences Research Council (BBSRC) grants BB/N005007/1 and BB/N005112/1 (as part of the ERA-CAPS consortium “MAQBAT”). HJS, RW, CNJ, GSS, RB, and CJR were also funded by the John Innes Institute strategic program grants Plant Health BB/P012574/1 (Response BBS/E/J/000PR9796), Molecules from Nature BB/P012523/1, Genes in the Environment BB/P013511/1 and Brassica Rapeseed and Vegetable Optimisation (BRAVO) BB/P003095/1.

## Author contributions

VKM did experiments with *A. brassicicola* infection and analysed the data with AKK. XZ did experiments and data curation with *V. longisporum* infection supervised by AvT. HF and GKM did experiments with *P. brassicae* and analysed data with HUS and HF, HUS and BDLF contributed to the conception and experimental design. RB and HJS did experiments and data curation with *B. cinerea* and PAMPs, HJS contributed to the conception and experimental design, and HJS and CNJ analysed the data. CNJ did statistical analysis of all datasets, CNJ and RW did association transcriptomics, GSS did the WGCNA analysis, and BS did the BLAST validation of the genomic deletion. CNJ, RW and CJR contributed to the conception, data interpretation and in-silico analyses post-AT. The first draft of the manuscript was written by CNJ, and all authors commented on subsequent versions of the manuscript. All authors read and approved the final manuscript.

## Data availability

The R-based GAGA pipeline is available at https://github.com/bsnichols/GAGA. Our population structure is available in Fell *et al*. (2022). Supplemental files are also available at https://zenodo.org/record/8321694.

