## Supporting Information for "Pathogen lifestyle determines host genetic signature of quantitative disease resistance loci in oilseed rape (*Brassica napus*)"

### *New Phytologist* Supporting Information

Article acceptance date: Click here to enter a date.

**The following Supporting Information is available for this article:**

**Fig. S1** Comparisons between chitin-, flg22- and elf18-induced ROS responses for the phenotypes and associated gene expression markers (GEMs).

**Fig. S2** Query coverage in *Brassica napus* reference genotypes and gene expression in *B. napus* population for region on chromosome C08 homoeologous to the region on A09 associated with resistance to *Verticillium longisporum*.

**Table S1** Mean, normalized phenotype data for resistance to pathogens (*Alternaria brassicicola, Botrytis cinerea, Pyrenopeziza brassicae* and *Verticillium longisporum*) and ROS response induced by PAMPS (chitin, flg22, and elf18).

**Table S2** Full list of single nucleotide polymorphism (SNP) markers and significance levels from genome-wide association (GWA) analyses for resistance to pathogens (*Alternaria brassicicola, Botrytis cinerea, Pyrenopeziza brassicae* and *Verticillium longisporum*) and ROS response induced by PAMPS (chitin, flg22, and elf18).

**Table S3** Full list of gene expression markers (GEMs) and significance levels from GEM analyses for resistance to pathogens (*Alternaria brassicicola, Botrytis cinerea, Pyrenopeziza brassicae* and *Verticillium longisporum*) and ROS response induced by PAMPS (chitin, flg22, and elf18).

**Table S4** 184 gene expression markers (GEMs) associated with chitin-induced ROS compared with GEMs associated with resistance to pathogens (*Alternaria brassicicola, Botrytis cinerea, Pyrenopeziza brassicae* and *Verticillium longisporum*) and ROS response induced by flg22, and elf18. Lists correspond to Venn diagrams in Fig. 2**.**

**Table S5** Enrichment analyses to determine if the number of gene expression markers (GEMs) shared between different lists is greater than the number of GEMs that would be expected by chance (e.g., lists of quantitative disease resistance (QDR) GEMs for two fungal pathogens).

**Table S6** Results from Weighted Co-expression Gene Network Analysis. Significant modules, significant GEM markers within modules, and GO terms associated with the magenta and black modules are indicated.

**Table S7** Shared gene expression markers (GEMs) associated with resistance to different pathogens (*Alternaria brassicicola, Botrytis cinerea, Pyrenopeziza brassicae* and *Verticillium longisporum*). Lists correspond to matrices and Venn diagrams in Fig. 1.

**Table S8** List of genes in linkage disequilibrium with the top marker for *Verticillium longisporum* resistance from genome-wide association (GWA) analysis on chromosome A09, the homoeologous region on C08, and their query coverage in *Brassica napus* reference genotypes

**Fig. S1 There is a correlation between phenotypes and GEMs associated with chitin-, flg22-, and elf18-induced ROS** (**a**) Flg22- and elf18-induced ROS response (arbitrary units (a.u.) normalized values between zero and one) of different *B. napus* genotypes. The position of reference genotypes Quinta (Q), Tapidor (T), Westar (W) and Zhongshuang 11 (Z) are indicated. (**b**) Correlation between the PAMP-induced ROS responses of *B. napus* lines elicited by chitin, flg22, and elf18. Positive correlations (green) are indicated between pair-wise comparisons of ROS responses. (**c**) Venn diagrams showing the overlap between GEMs associated with chitin-, flg22-and elf18-induced ROS.

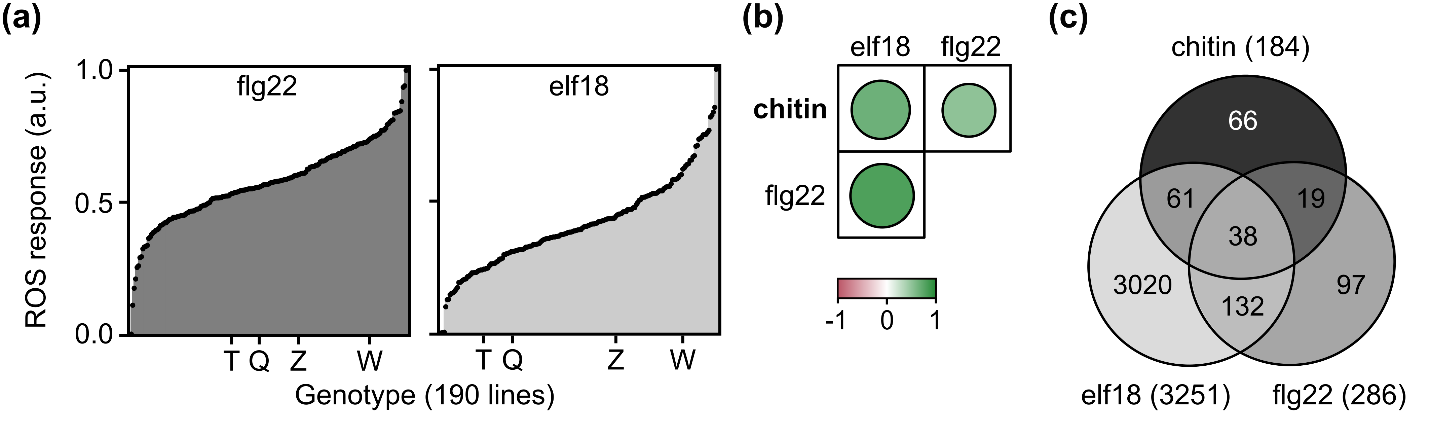

**Fig. S2 Chromosome C08 region homologous to the region on A09 associated with resistance to *Verticillium longisporum.*** (a) Heatmap indicating query coverage in Quinta (Q), Tapidor (T), Westar (W) and Zhongshuang 11 (Z) (compared to the *Brassica napus* pantranscriptome) for a region on chromosome C08, homoeologous to the region on A09 containing 107 genes predicted to be in linkage disequilibrium with Cab013526.1.651.A. The C08 homeolog, *Bo8g101810.1*, of *Cab013526.1* (A09) is indicated. (b) Linear regression analysis of gene expression (RPKM) of the five Gene Expression Markers (GEMs) in the genomic deletion on chromosome A09 and their homeologs on chromosome C08 relative to resistance to V*. longisporum* (arbitrary units (a.u.) normalized values between zero and one) in 191 *B. napus* genotypes

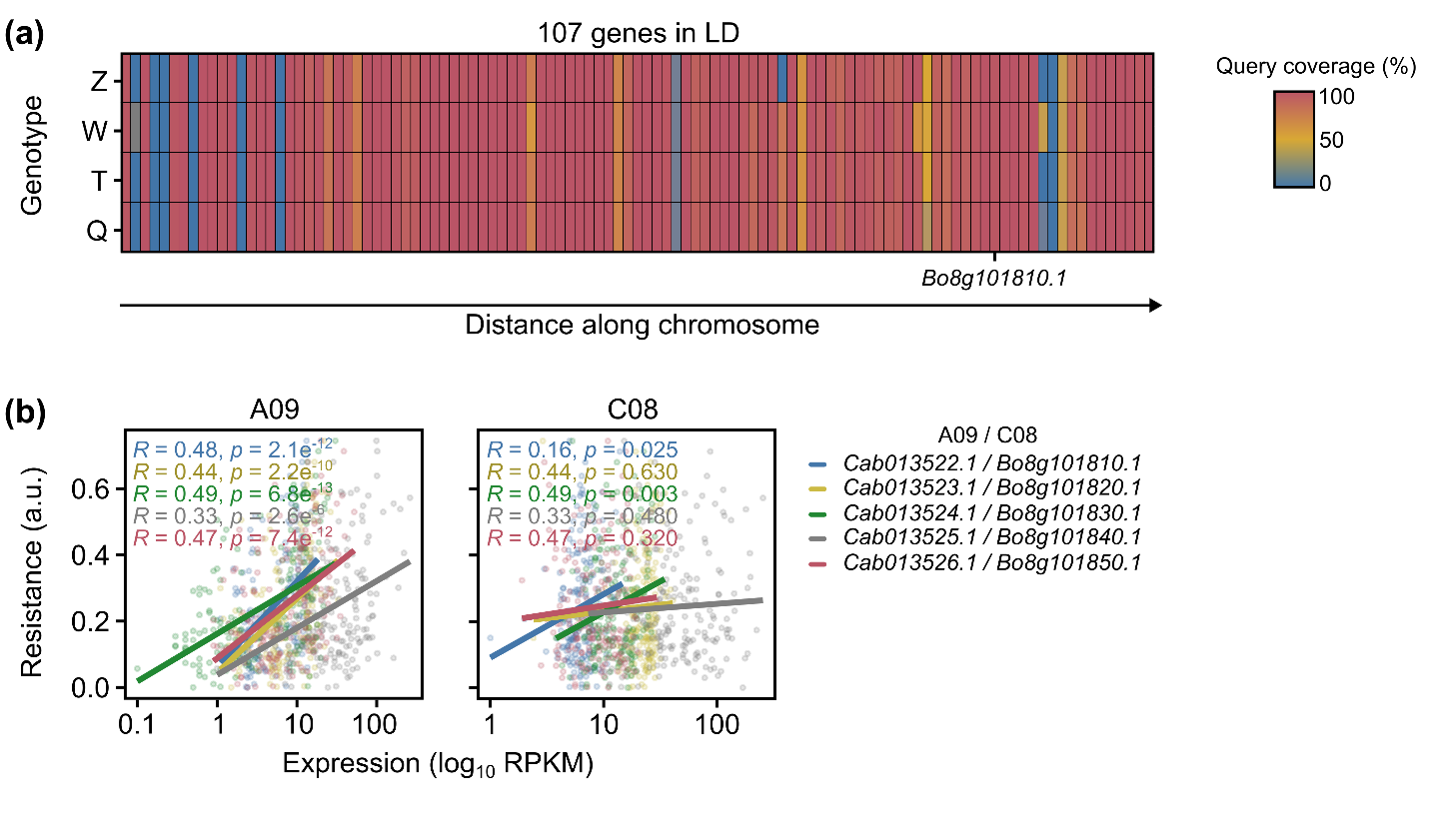

**Table S1** Mean, normalized phenotype data for resistance to pathogens (*Alternaria brassicicola, Botrytis cinerea, Pyrenopeziza brassicae* and *Verticillium longisporum*) and ROS response induced by PAMPS (chitin, flg22, and elf18). These data were used for association transcriptomic analysis. **Excel file separate**

**Table S2** Full list of single nucleotide polymorphism (SNP) markers and significance levels from genome-wide association (GWA) analyses for resistance to pathogens (*Alternaria brassicicola, Botrytis cinerea, Pyrenopeziza brassicae* and *Verticillium longisporum*) and ROS response induced by PAMPS (chitin, flg22, and elf18). Each excel tab contains the analyses for a single trait. The best fit model for GWA analysis is indicated in the tab title. Manhattan plots showing marker-trait association are included for data visualization; x-axis indicates SNP location along the chromosome; the y-axis indicates the -log10(p) (P value). Qqplots are included to demonstrate model fit. **Excel file separate**

**Table S3** Full list of gene expression markers (GEMs) and significance levels from GEM analyses for resistance to pathogens (*Alternaria brassicicola, Botrytis cinerea, Pyrenopeziza brassicae and Verticillium longisporum*) and ROS response induced by PAMPS (chitin, flg22, and elf18). Each excel tab contains the analyses for a single trait. Manhattan plots showing marker-trait association are included for data visualization; x-axis indicates GEM location along the chromosome; the y-axis indicates the -log10(p) (P value). **Excel file separate**

**Table S4** 184 gene expression markers (GEMs) associated with chitin-induced ROS compared with GEMs associated with resistance to pathogens (*Alternaria brassicicola, Botrytis cinerea, Pyrenopeziza brassicae* and *Verticillium longisporum*) and ROS response induced by flg22, and elf18. Lists correspond to Venn diagrams in Fig. 2. The first tab includes all 184 GEMs associated with chitin-induced ROS. The subsequent tabs include lists of shared GEMs associated with chitin-induced ROS response and each additional trait (quantitative disease resistance (QDR) to each fungal pathogen or additional PAMP-induced ROS responses). The title of each tab indicates the data included in each comparison and the number of shared GEMs. Predicted *Arabidopsis thaliana* orthologs and corresponding descriptions are shown where possible. **Excel file separate**

**Table S5** Enrichment analyses to determine if the number of gene expression markers (GEMs) shared between different lists is greater than the number of GEMs that would be expected by chance (e.g., lists of quantitative disease resistance (QDR) GEMs for two fungal pathogens). The representation factor is the number of overlapping GEMs divided by the expected number of overlapping GEMs drawn from two independent groups (traits), considering the total number of GEMs sequenced (53884). A representation factor > 1 indicates more overlap than expected of two groups, a representation factor < 1 indicates less overlap than expected, and a representation factor of 1 indicates that the two groups by the number of genes expected for independent groups of genes. **Excel file separate**

**Table S6** Results from Weighted Co-expression Gene Network Analysis (WGCNA). The first tab indicates significant modules from WGCNA analysis. Black and magenta modules are associated with antagonistic effects on resistance/susceptibility to all four pathogens. The second tab includes a full list of the GEM markers (Table S3), which are in significant WGCNA modules. The third, fourth and, fifth tabs indicate all significant GEMs in the black module, GO terms associated with GEMs in the black module, and all GO terms associated with the black module, respectively. The sixth, seventh and, eighth tabs indicate all significant GEMs in the magenta module, GO terms associated with GEMs in the magenta module, and all GO terms associated with the magenta module, respectively. **Excel file separate**

**Table S7** Shared gene expression markers (GEMs) associated with resistance to different pathogens (*Alternaria brassicicola, Botrytis cinerea, Pyrenopeziza brassicae* and *Verticillium longisporum*). Lists correspond to matrices and Venn diagrams in Fig. 3. The first tab includes all GEMs associated quantitative disease resistance (QDR) to the fungal pathogens. The subsequent tabs include lists of shared GEMs associated with QDR to two or more fungal pathogens. The title of each tab indicates the data included in each comparison and the number of shared GEMs. Predicted *Arabidopsis thaliana* orthologs and corresponding descriptions are shown where possible. **Excel file separate**

**Table S8** List of genes in linkage disequilibrium with the top marker for *Verticillium longisporum* resistance from genome-wide association (GWA) analysis on chromosome A09 (107 genes)(Tab 1) and the homoeologous region on C08 (Tab 2). Their percentage identity and query coverage in *Brassica napus* reference genotypes Quinta, Tapidor, Westar and Zhongshuang 11 compared to the *B. napus* pantranscriptome is indicated. Predicted *Arabidopsis thaliana* orthologs and corresponding descriptions are shown where possible. **Excel file separate**
